# Assessment of Genetic Diversity and Population Genetic Structure of *Santalum album* L. in India by Genic and Genomic SSR markers

**DOI:** 10.1101/2021.05.28.446175

**Authors:** Tanzeem Fatima, Ashutosh Srivastava, Vageeshbabu S. Hanur, M. Srinivasa Rao

## Abstract

Sandalwood (*Santalum album* L.) is highly valued aromatic tropical tree. It is known for its high quality heartwood and oil. In this study, 39 genic and genomic SSR markers were used to analyze the genetic diversity and population structure of 177 *S. album* accessions from 14 populations of three states in India. High genetic diversity was observed in terms of number of alleles 127, expected heterozygosity (He) ranged from 0.63-0.87 and the average PIC was 0.85. The selected population had relatively high genetic diversity with Shannon’s information index (I) >1.0. 0.02 mean coefficient of genetic differentiation (F_ST_) and 10.55 gene flow were observed. AMOVA revealed that 92% of the variation observed within individuals. Based on cluster and Structure result, individuals were not clustered as per their geographical origin. Furthermore the clusters were clearly distinguished by principal component analysis analysis and the result revealed that PC1 reflected the moderate contribution in genetic variation (6%) followed by PC2 (5.5%). From this study, high genetic diversity and genetic differentiation was found in *S. album* populations. The genetic diversity information of *S. album* populations can be used for selection of superior genotypes and germplasm conservation to promote the tree improvement of *S. album* populations.

## 1. Introduction

Indian Sandalwood (*Santalum album* L.; Family-Santalaceae) is an economically important tropical tree species (Kuriakose et al., 2010). It is naturally distributed in India, Indonesia, Australia, Sri Lanka, Fiji, pacific islands, Juan Fernandes islands, Niue and Tonga (Harbaugh and Baldowin, 2007; Subasinghe, 2013). India has approximate area of 9000 sq. kms of *S. album* distribution spread across eight states; Karnataka having the maximum area of 5, 245 sq. kms, Tamil Nadu (3,040 sq. kms), Andhra Pradesh (200 sq. kms), Kerala (15 sq. kms), Odisha (25 sq. kms), Madhya Pradesh (33 sq. kms) and Maharashtra (33 sq. kms) (Kumaravelu et al., 2007). It is also distributed in other states, Assam, Rajasthan, Uttar Pradesh, Bihar and Manipur (Ananthapadmanabha and Gupta 2016). The scented heartwood and its essential oil served for religious purposes, sophisticated perfumes, flavors, cosmetics, toiletries, beauty aids and medicines (Sreenivasan et al., 1992). It has a great demand in traditional Ayurveda medicines for antidepressant, anti-inflammatory, anti-fungal, astringent, sedative, as an insecticide, antipyretic and antiseptic (Kirthikar and Basu, 1987; Heubeger et al., 2006; Burdock et al., 2008; Silva et al., 2018). Due to high rate of cross-breeding and the extensive range of agro-climatic conditions, *S. album* prevailed high rate of genetic diversity and complex genetic structure (Veerendra and Anathapadmanabha, 1996). In earlier studies it was reported that a large gap in genetic distance of *S. album* populations, belongs to different geographical regions (Angadi et al., 2003). Various of factors including illicit felling, large-scale changes in land use and poor natural regeneration, superior genetic resources of *S. album* in the country are severely threatened and natural populations have been depleted considerably (Sreenivasan *et al*., 1992; Rao et al., 2007; Jain et al., 2003). Due to amendment to the Karnataka Forest Act 2001, many of the private organizations and people have started raising plantations. Under this scenario it is essential to assess the morphological and genetic diversity of *S. album* natural populations and plantations. Although the morphological parameter reflects the diversity they are mostly influenced by environmental factors. Therefore molecular markers are being extensively used to assess genetic diversity of the species. To date various dominant and co-dominant marker system has been used in genetic diversity and population structure study of *S. album* (RFLP, Isozymes, DAMD, Allozymes, and RAPD) markers (Brand 1994; Suma and Balasundaran, 2003; Shashidhara et al., 2003; Rao et al., 2007; Jones et al., 2008; Azeez et al., 2009; Dani et al., 2011). Microsatellite SSR markers genic and genomic have proven to be efficient and accurate tool for the study of genetic variation in forest tree species compared with other markers (Kalia et al., 2011). However, very few genomic and genic microsatellite markers have been developed for *S. album* (Naseer et al., 2012; Patel et al. 2016; Fatima et al., 2019a) to characterize genetic diversity and genetic differentiation of *S. album*. Therefore in this study, we aimed to evaluate the genetic diversity and reveal the population structure of the natural populations and plantations of *S. album* by using both genomic and genic microsatellite markers. These results facilitate novel insight into the genetic diversity and population genetic structure in *S. album* with more appropriateness. Evaluation of genetic diversity could useful in tree improvement, tree breeding process and conservation management of *S. album* populations.

## 2. Material and Methods

The basic objective of the current study was to assess genetic diversity of n = 177 *S. album* accessions from natural populations and plantations of three southern states in India *viz*, Karnataka (n=80), Kerala (n=30) and Telangana (n=67) (Table 1). Genetic diversity analysis of *S. album* was performed by 39 (14 genic and 25 random genomic SSR markers). The DNA isolation, PCR and gel electrophoresis details were available (Fatima et al., 2019a & Fatima et al., 2019b).

**Table 1.**
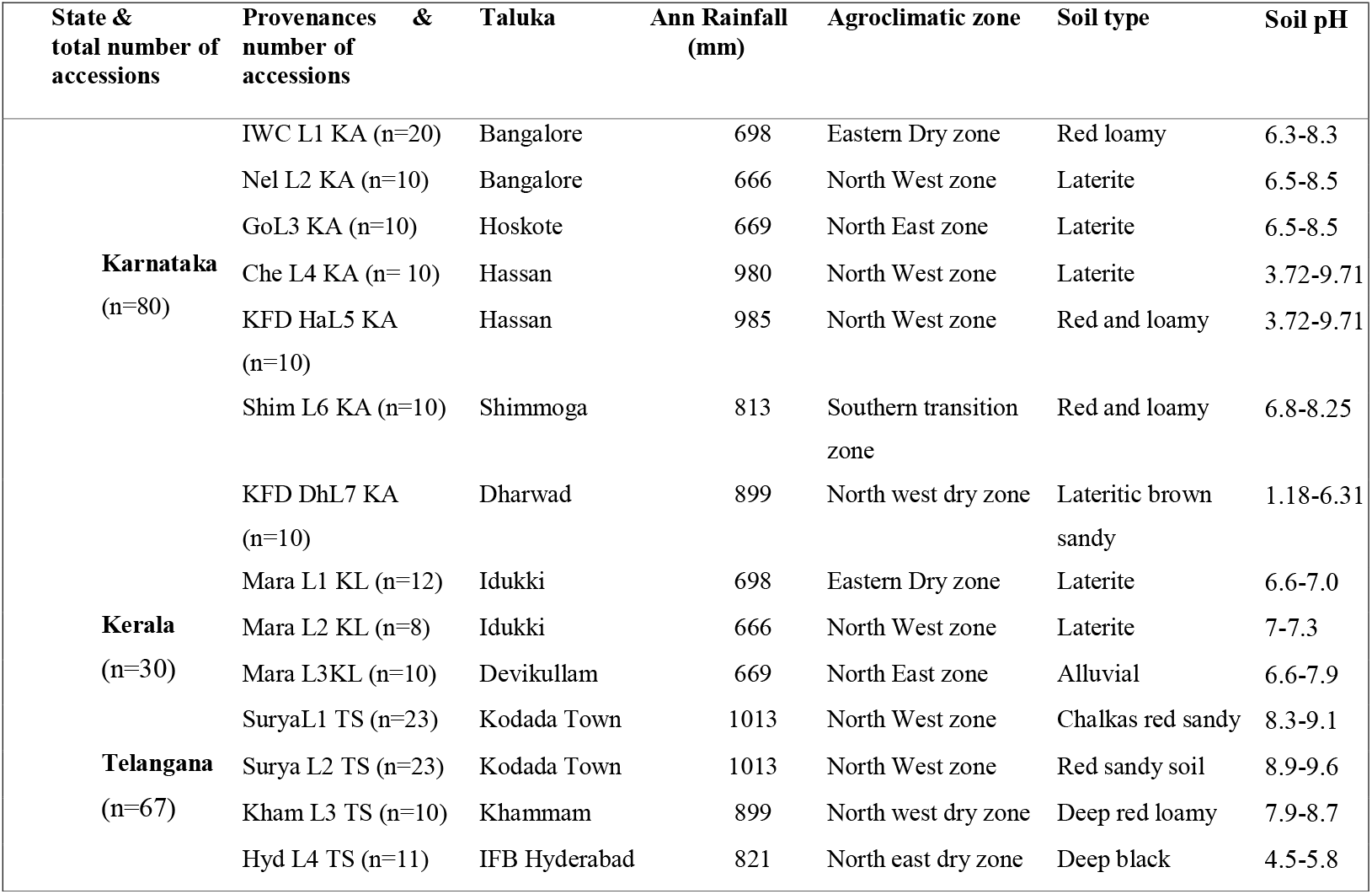
Details of *S. album* natural populations and plantations identified across Karnataka, Kerala and Telangana state

### 2.1. Genetic Diversity Parameters, Genetic Differentiation, Cluster analysis

Genetic diversity parameters; number of alleles (Na), effective number of allele (Ne), observed heterozygosity (Ho), expected heterozygosity (He), Shannon information index (I), Nei’s genetic distance and Polymorphic Information Content (PIC) were characterized by GenAlex 6.0 software and Cervus 3.0.3 software (Peakall and Smouse, 2012; Kalinowski et al., 2007). To determine the genetic differentiation, analysis of molecular variance (AMOVA) and Wright’s *F* statistics were performed by GenAlex v.6.5 (Peakall and Smouse, 2012). Phylogenetic cluster analysis based on the UPGMA was constructed by Jaccard’s similarity coefficient by using Darwin software (version 6.0) (Takezaki et al., 2014) at 0.05 significance level.

### 2.2. Structure and Principal Component Analysis (PCA)

For analyzing population structure, a model-based (Bayesian) cluster analysis was performed based on SSR markers. An unbiased Bayesian approach using Markov chain Monte Carlo (MCMC) clustering of samples was conducted via the STRUCTURE v.2.3.4 software (http://www.stats.ox.ac.uk/) (Pritchard et al. 2000). PCA was executed for selected *S. album* accessions data by using Minitab v.18 software (McKenzie, 2004). It was used to identify the similarities between variables and classify the genotypes collections from different geographical locations (Torbick and Becker, 2009).

## 3. Results

### 3.1. Population genetic diversity and variation

In the current study, 127 alleles were amplified by 39 SSR loci examined in the 177 *S. album* accessions and the average Na detected 9.39 (Table 2). Among the selected populations the largest Na (12.13), Ne (9.12), He (0.87) was harbored in Telangana population Surya L2 TS (Table 2). The highest I 2.24 and PIC (0.96) was observed in Karnataka population IWC L1 KA respectively (Table 2). However, the Karnataka populations Chen L4KA showed lower than average Ne (7.90) and He (0.83) (Table 2). The lowest PIC was found in Kerala population Mara L3 KL (Table 2). In terms of PIC value, 10 populations (WC L1KA, NelL2KA, KFDHaL5KA, KFDDhL7 KA, ShimL6KA, GoL KA, Surya L2 TS, Kham L3 TS and HydL4TS) had highly informative alleles with values higher than 0.80 while only population (ChenL4KA, MaraL1 KA, MaraL2 KL, Mara L3 KL) had moderately informative alleles with a PIC value between 0.72-0.79 (Table 2). The average F, F_ST_ and Nm 0.80, 0.02 and 10.55 respectively (Table 2 and Table 4).

**Table 2.**
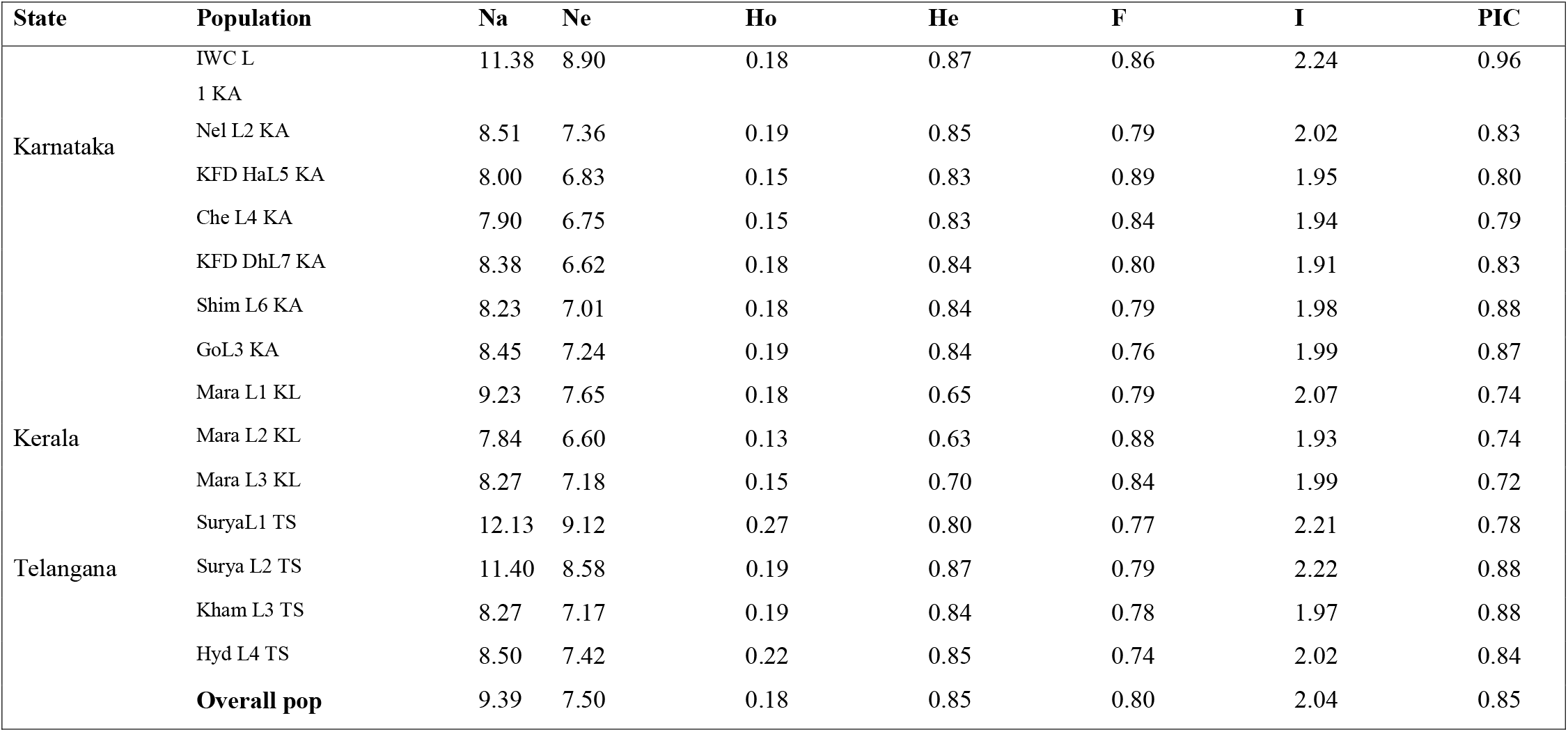
Number of alleles (Na), effective number of alleles (Ne), observed heterozygosity (Ho), Expected heterozygosity (He), fixation index (F), Polymorphic Information Content (PIC) of fourteen populations of *S. album* by 39 genic and genomic SSR markers

### 3.2. Population genetic structure and genetic differentiation

Both AMOVA and pairwise F_ST_ analysis were conducted to investigate the genetic variation among 14 populations. Hierarchical AMOVA revealed that total 3% of the total genetic variation occurred among 14 populations and 92% of the total variation was distributed within the individuals (Table 3). Population genetic structure of *S. album* revealed strong population genetic structure at the species level F_ST_ (0.08, P<0.001) (Table 3). AMOVA indicated that a small fraction of the observed genetic diversity was attributable to differences among the populations (Table 3).

**Table 3.**
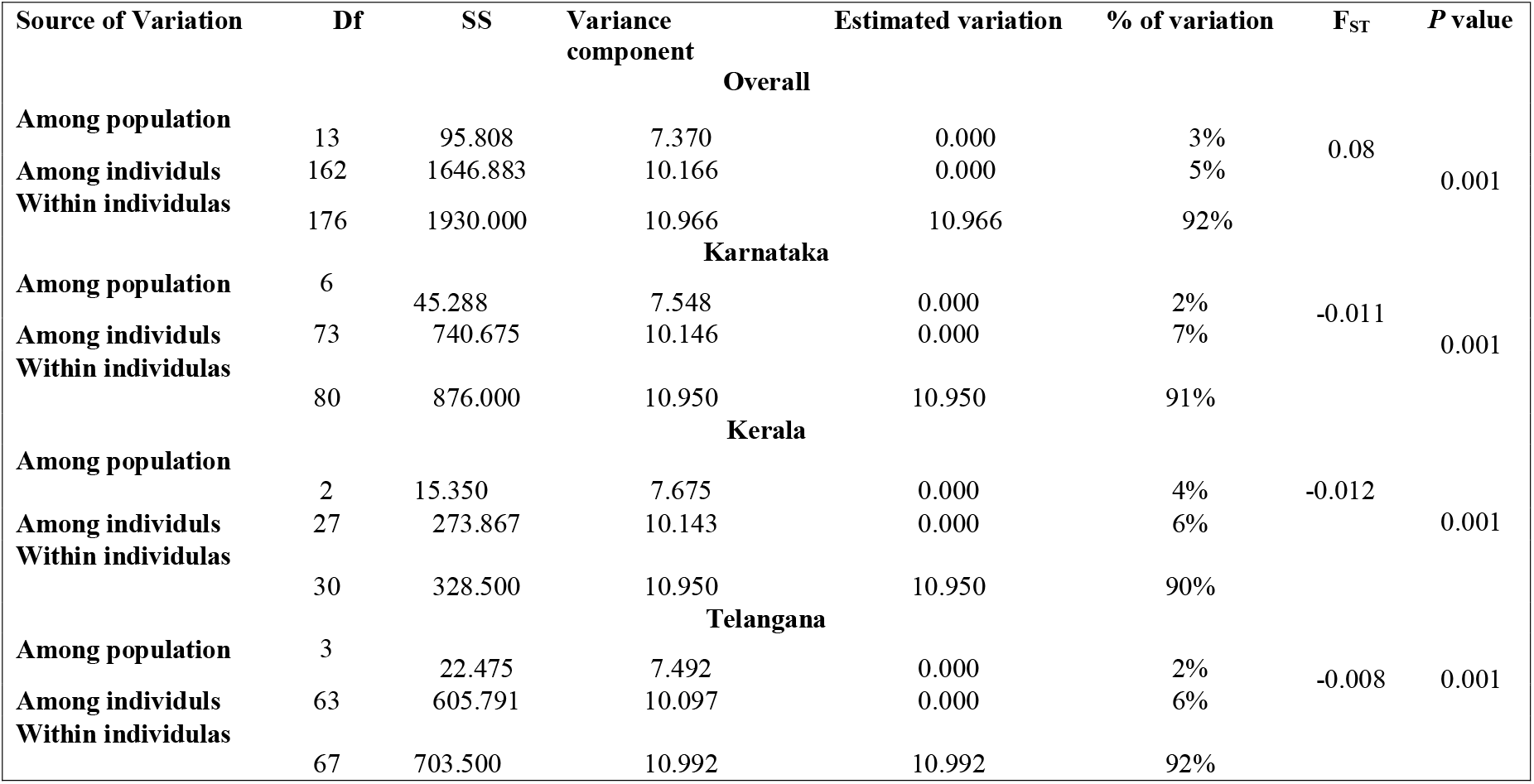
Analysis of Molecular Variance (AMOVA) of 177 *S. album* accesssions based on 39 genic and genomic SSR markers

**Table 4.** F statistics and Nm (Gene flow) of 14 populations of *S. album* by 39 genic and genomic SSR markers; Nm = [(1/Fst)-1]/4

| **States** | **Populations** | **F<sub>ST</sub>** | **Nm** |
| --- | --- | --- | --- |
| **Karnataka** | IWC L1 KA | 0.03 | 9.50 |
|  | Nel L2 KA | 0.03 | 8.18 |
|  | KFD DhL7 KA | 0.02 | 12.95 |
|  | KFD HaL5 KA | 0.020 | 10.75 |
|  | Che L4 KA | 0.02 | 8.75 |
|  | GoL3 KA | 0.02 | 9.57 |
|  | Shim L6 KA | 0.02 | 8.06 |
| **Kerala** | Mara L1 KL | 0.02 | 13.47 |
|  | Mara L2 KL | 0.02 | 8.66 |
|  | Mara L3 KL | 0.02 | 8.66 |
| **Telangana** | SuryaL1 TS | 0.02 | 14.57 |
|  | Surya L2 TS | 0.02 | 12.44 |
|  | Kham L3 TS | 0.03 | 10.51 |
|  | Hyd L4 TS | 0.02 | 11.62 |
| **Average** |  | 0.02 | 10.55 |

Cluster analysis based on the neighbour-joining (NJ) dendrogram method was established to demonstrate the relationship between 177 accessions of *S. album* from 14 populations was clearly clustered into four clusters. Cluster I comprised three natural populations from Karnataka and Kerala *viz*, Mara L3KL, Shim L6 KA, and Che L4 KA. Cluster II consisted of 11 natural populations and plantations admixtures *viz*, SuryaL1 TS, Surya L2 TS, Mara L1 KL, Mara L2 KL, Hyd L4 TS, KFD DhL7 KA, IWC L1 KA, Nel L2 KA, Kham L3 TS, KFD HaL5 KA, and GoL3 KA. In cluster I natural populations and plantations both were grouped together whereas, cluster II only plantations were segregated. Notably many neighboring populations cluster together specifically all four populations of Telangana state cluster II (Fig. 1). The pairwise population matrix between any two provenances, the index values varied from 0.04-0.286. The smallest value 0.04 was found in between IWCL1 KA and ChenL4 KA along with the highest distant differentiation (0.286) was observed in Mara L2 KL and KFD DhL7 KA, which originated from two geographical distant states in southern India (Table 5).

**Fig. 1.**
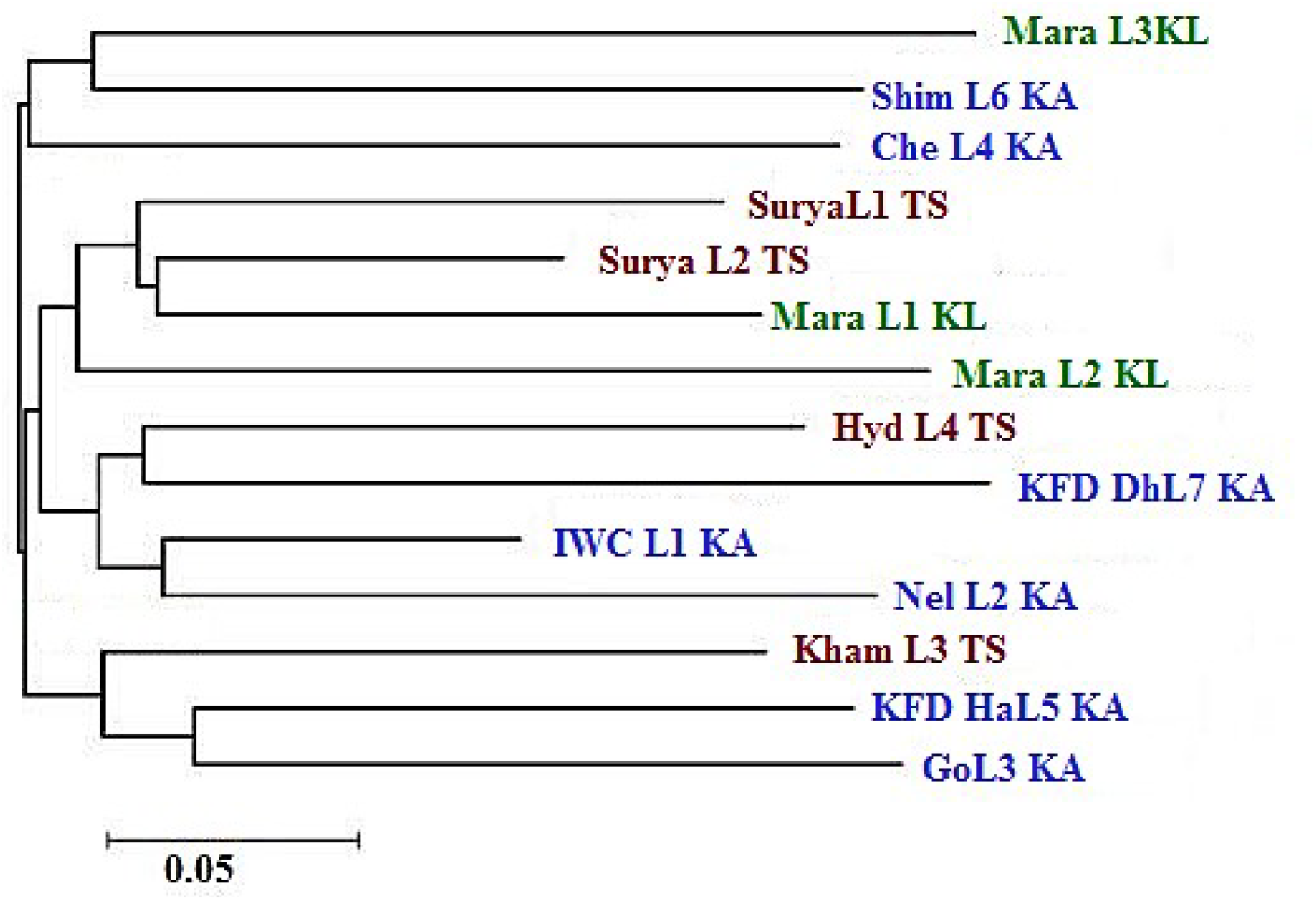
Neighbour joining tree based UPGMA dendrogram showing the clustering pattern of 14 populations of *S. album* using 39 microsatellite markers (*Red:Telangana population; Blue: Karnataka populations; Green: Kerala populations).

**Table 5.** Pairwise Population Matrix of Nie’s (1973) unbiased genetic distance by using 39 genic and genomic SSR markers

| Surya L1 TS | Surya L2 TS | Kham L3 TS | Hyd L4 TS | Mara L1 KL | Mara L2 KL | Mara L3 KL | IWC L1 KA | Nel L2 KA | KFD HaL5 KA | Che L4 KA | KFD DhL7 KA | Shim L6 KA | GoL3 KA |  |
| --- | --- | --- | --- | --- | --- | --- | --- | --- | --- | --- | --- | --- | --- | --- |
| *** |  |  |  |  |  |  |  |  |  |  |  |  |  | SuryaL1 TS |
| 0.08 | *** |  |  |  |  |  |  |  |  |  |  |  |  | SuryaL2 TS |
| 0.09 | 0.01 | *** |  |  |  |  |  |  |  |  |  |  |  | Kham L3 TS |
| 0.08 | 0.06 | 0.04 | *** |  |  |  |  |  |  |  |  |  |  | Hyd L4 TS |
| 0.04 | 0.00 | 0.00 | 0.06 | *** |  |  |  |  |  |  |  |  |  | Mara L1 KL |
| 0.14 | 0.12 | 0.08 | 0.00 | 0.14 | *** |  |  |  |  |  |  |  |  | Mara L2 KL |
| 0.19 | 0.09 | 0.11 | 0.05 | 0.09 | 0.193 | *** |  |  |  |  |  |  |  | Mara L3 KL |
| 0.09 | 0.05 | 0.01 | 0.00 | 0.00 | 0.098 | 0.11 | *** |  |  |  |  |  |  | IWC L1 KA |
| 0.18 | 0.10 | 0.05 | 0.00 | 0.11 | 0.078 | 0.09 | 0.07 | *** |  |  |  |  |  | Nel L2 KA |
| 0.12 | 0.01 | 0.00 | 0.03 | 0.07 | 0.123 | 0.16 | 0.00 | 0.19 | *** |  |  |  |  | KFD HaL5 KA |
| 0.07 | 0.09 | 0.03 | 0.00 | 0.05 | 0.064 | 0.16 | 0.04 | 0.13 | 0.05 | *** |  |  |  | Che L4 KA |
| 0.22 | 0.18 | 0.07 | 0.12 | 0.17 | 0.286 | 0.17 | 0.10 | 0.17 | 0.05 | 0.14 | *** |  |  | KFD DhL7 KA |
| 0.14 | 0.05 | 0.09 | 0.03 | 0.03 | 0.153 | 0.13 | 0.02 | 0.04 | 0.09 | 0.15 | 0.22 | *** |  | Shim L6 KA |
| 0.14 | 0.10 | 0.12 | 0.10 | 0.13 | 0.171 | 0.08 | 0.04 | 0.16 | 0.08 | 0.08 | 0.13 | 0.13 | *** | GoL3 KA |

Structure analysis with the proportion of cluster members of each accession from 14 populations is shown in (Fig. 2). Individuals with a proportion higher than 0.75 were considered pure and those with lower 0.75 were considered as admixtures. The graph showed that the geographical distance was relatively not clear. The first cluster (red lines) had a lower proportion than second cluster (green lines) (Fig. 2). Structure analysis was conducted for highest value K = 4 which means that all populations were divided into four genetic structure groups with admixtures. At K=4 the estimated Ln probability of all data was <0.75, inferred cluster value 28.85% (red lines) 30.45% (green lines) 20.34% blue and 22.45% (yellow lines) revealed that genetic structure of all the selected populations showed admixtures of mostly in Telangana and Karnataka state accessions whereas, Kerala populations were found pure showed in yellow lines (Fig. 2). State wise, structure analysis and cluster analysis of all accessions (n=80) from Karnataka revealed four clusters (ΔK=4). The lines were divided into two groups I, II III and IV groups with admixtures. The mean assignment probabilities were <0.7 **i.** (G1 18) (red) with the dendrogram individuals group I and **ii.** (G2 31) (green) **iii.** (G3 16) (blue) and **iv.** (G4 15) (yellow) individuals were observed (Fig. 3). Similarly in Kerala populations revealed four major groups (ΔK=4) and the accessions were divided into four groups I, II, III and IV with the mean assignment probabilities <0.8 **i.** (G1 11) (red) **ii.** (G2 9) (green) and **iii.** (G3 4) (blue) and **iv.** (G4 8) (yellow) (Fig. 4). Structure analysis of four populations of Telangana with (n=65) accessions divided into three major groups at Δ K= 3. The mean assignment probabilities at <0.9 inferred cluster value **i.** (G1 36) (red) **ii.** (G2 19) (green) **iii.** (G3 10) (blue). Around 16 accessions from Kodada, Suryapet and IFB Hyderabad was found admixtures (Fig. 5). This result was consistent with that of UPGMA clustering analysis (Fig. 1, 2, 3, 4 and 5). Similarly the three clusters were clearly distinguished by PCA analysis (Fig. 6); the populations from Karnataka, Kerala and Telangana were mixed and scattered in all three clusters (Fig. 6). The localization of individuals was defined by the first principal component (PC1) and second principal component (PC2). The results revealed that PC1 reflected the highest contribution in genetic variation (6%) followed by PC2 (5.5%). Eigen values of 25 PC components were found higher than 1 and 15 PC components were found to be lower than 1 revealed high variability in selected *S. album* accessions. The result showed the combined variation 87.5% and cumulative variation 98.6% (Table 6). The obtained result showed that there was no significant correlation was found between genetic distance and geographic distance among the investigated populations.

**Fig. 2.**
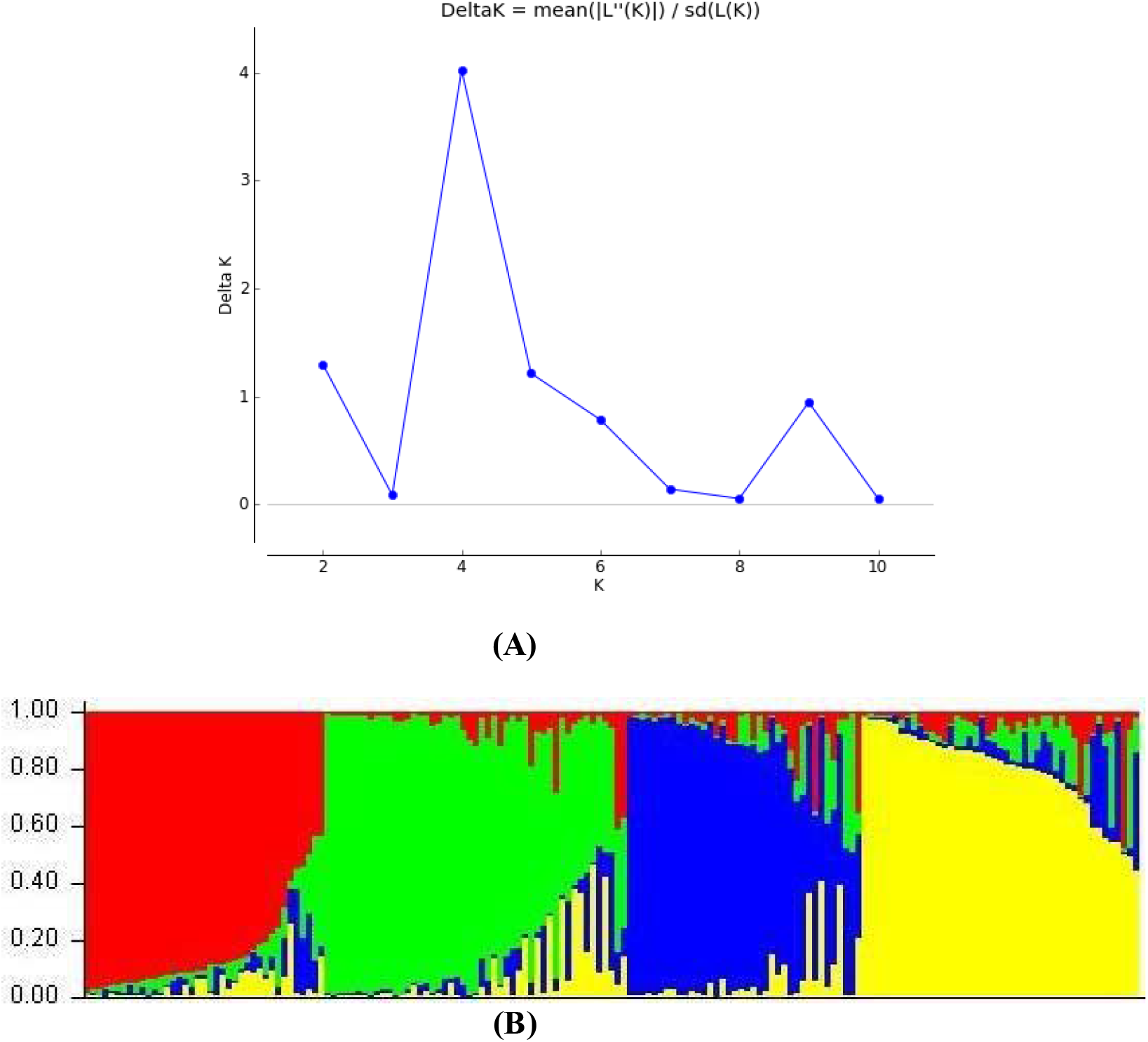
Result of Bayesian structure analysis for 177 *S. album* accessions based on 39 microsatellite markers **(A)** Delta K values from the mean log-likelihood probabilities from STRUCTURE runs with inferred cluster (K) ranged from 1-11 **(B)** Distribution of cluster memberships at the individual levels estimation using STRUCTURE software (Red area: Group I), (Green area: Group II), (Blue area: Group III), (Yellow area: Group IV).

**Fig. 3.**
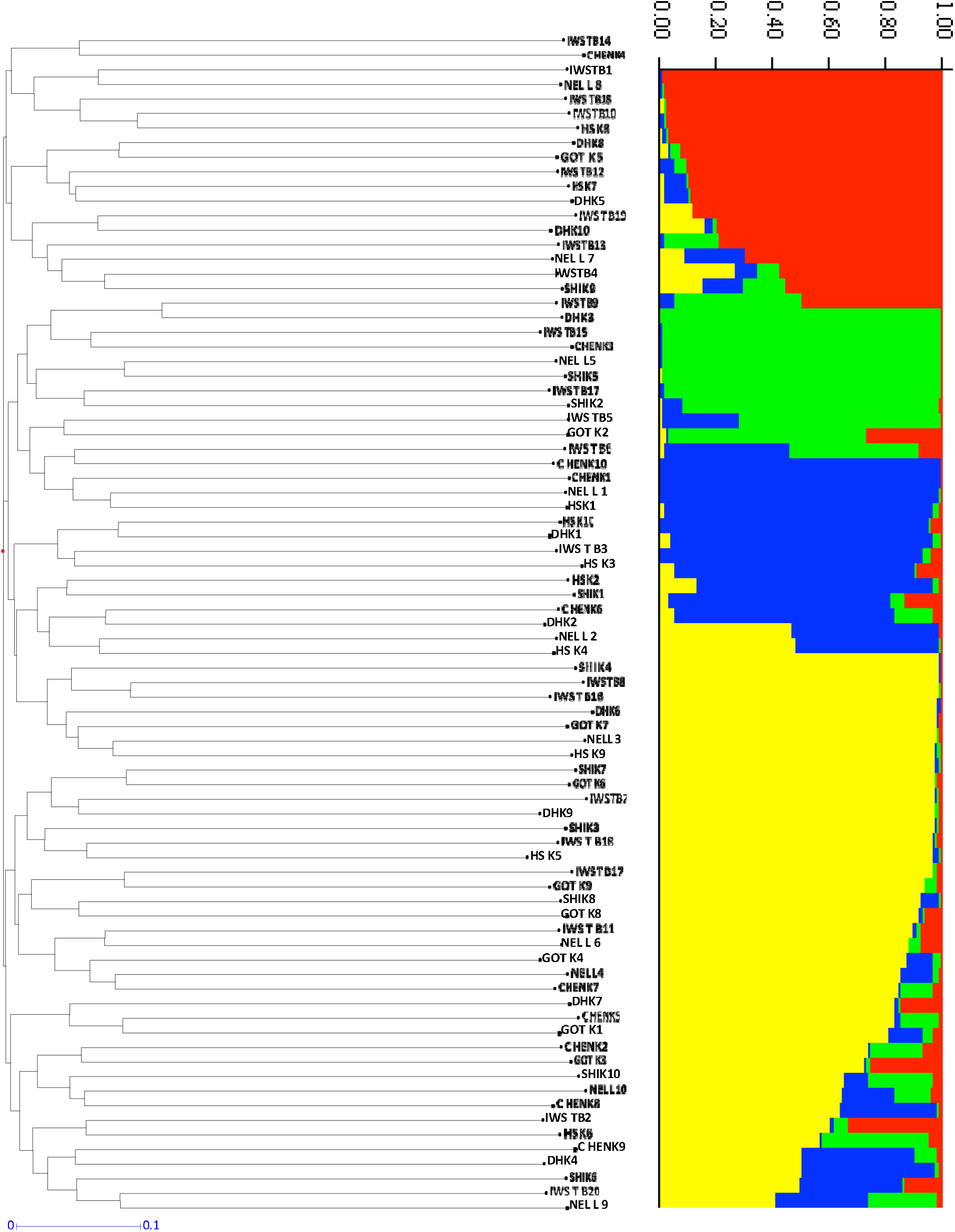
Genetic distance and structure analysis of *S. album* genotypes from 7 populations of Karnataka using 39 microsatellite markers **(ii)** The dendrogram constructed based on UPGMA analysis using similarity matrix **(ii)** The distribution of clustering of Population structure of Karnataka at estimated LnP (D) of possible clusters from (1-11) (K=4) Estimated genetic clustering (K=4) for 80 genotypes of Karnataka.

**Fig. 4.**
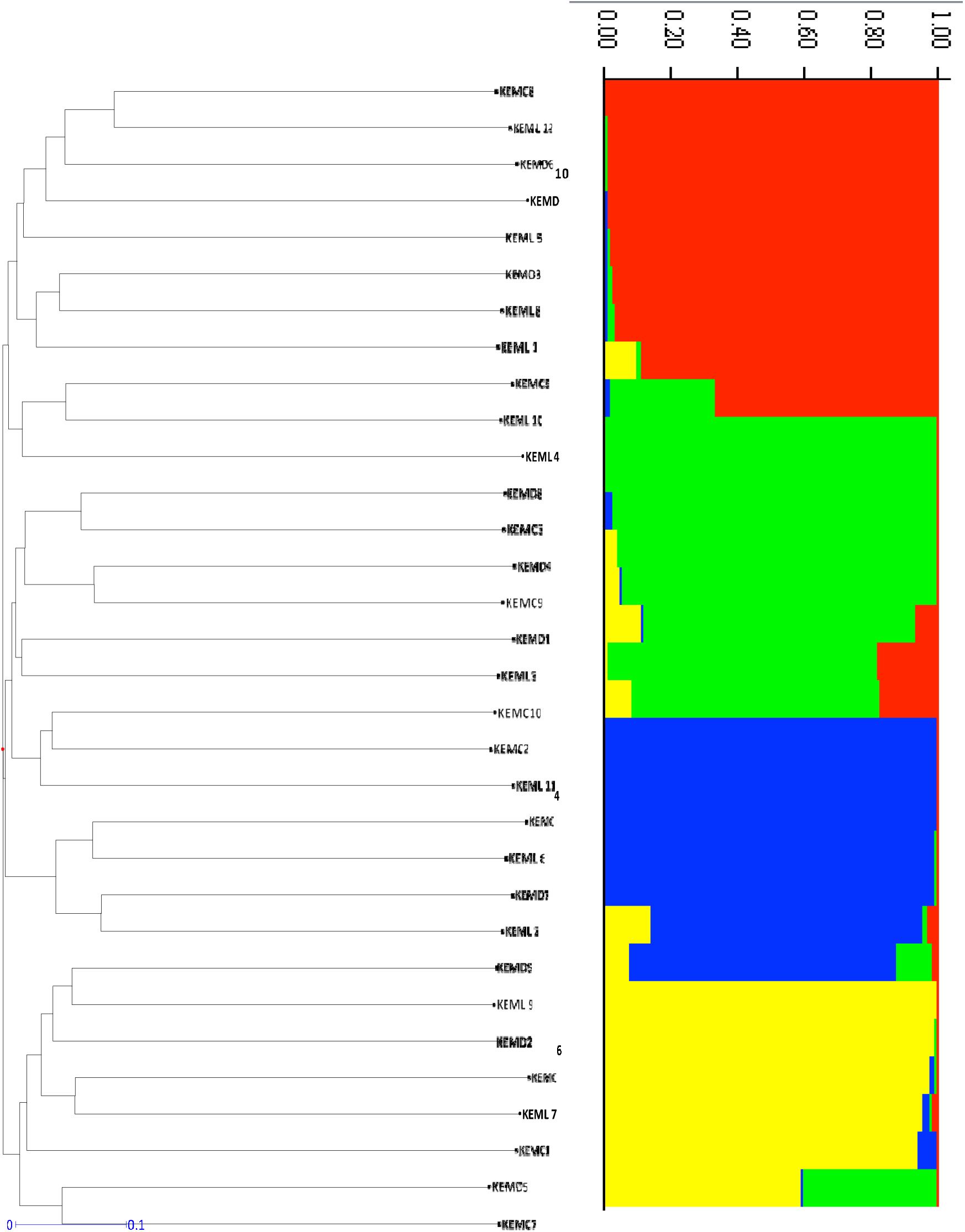
Genetic distance and structure analysis of *S. album* genotypes from 3 populations of Kerala using microsatellite markers **(i)** The dendrogram constructed based on UPGMA analysis using similarity matrix **(ii)** The distribution of clustering of Population structure of Karnataka at estimated LnP (D) of possible clusters from (1-11) (K=4) Estimated genetic clustering (K=4) for 30 genotypes of Kerala.

**Fig. 5.**
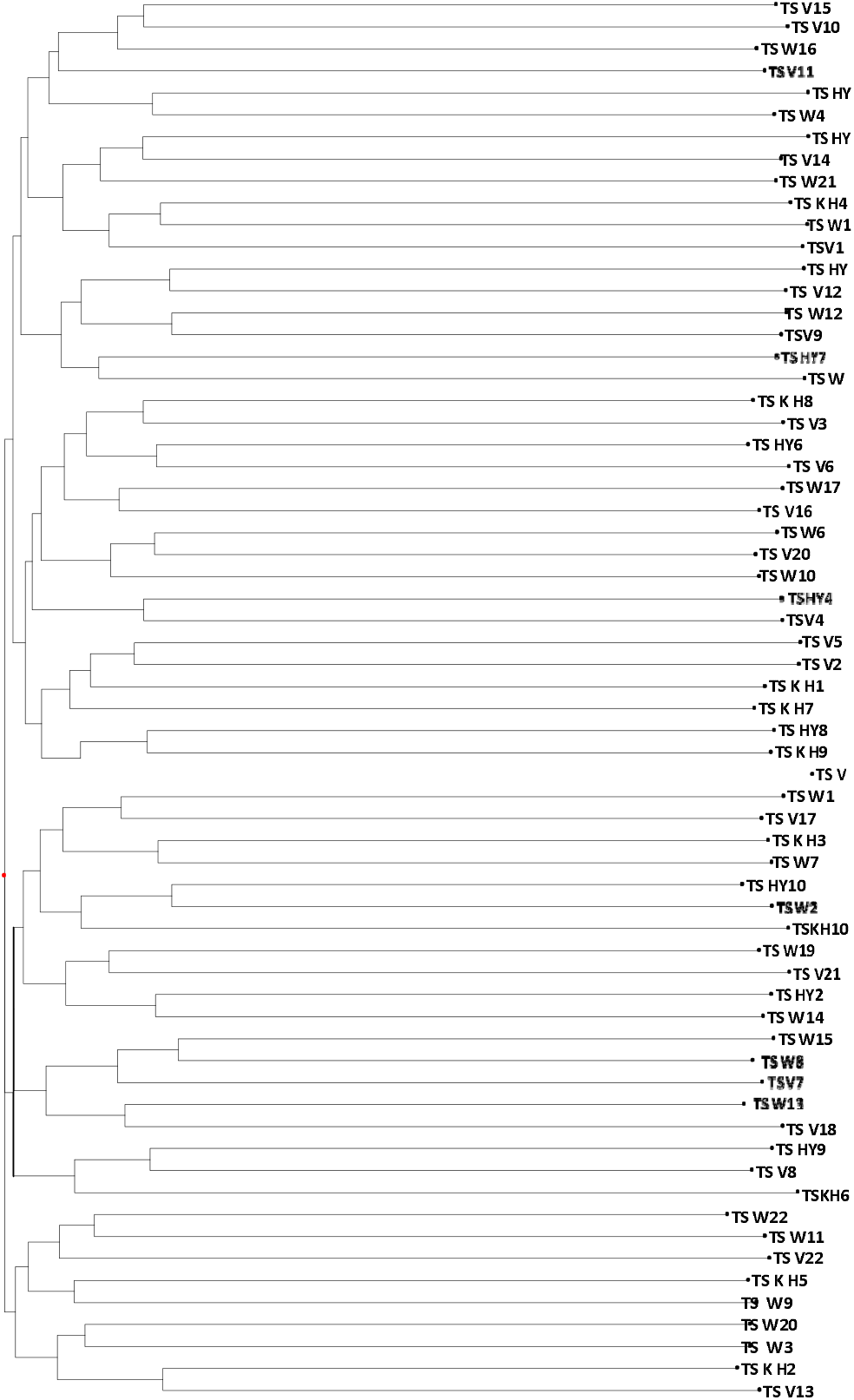
Genetic distance and structure analysis of *S. album* genotypes from 4 populations of Telangana using microsatellite markers **(i)** The dendrogram constructed based on UPGMA analysis using similarity matrix **(ii)** The distribution of clustering of Population structure of Karnataka at estimated LnP (D) of possible clusters from (1-11) (K=3) Estimated genetic clustering (K=4) for 64 genotypes of Telangana

**Fig. 6.**
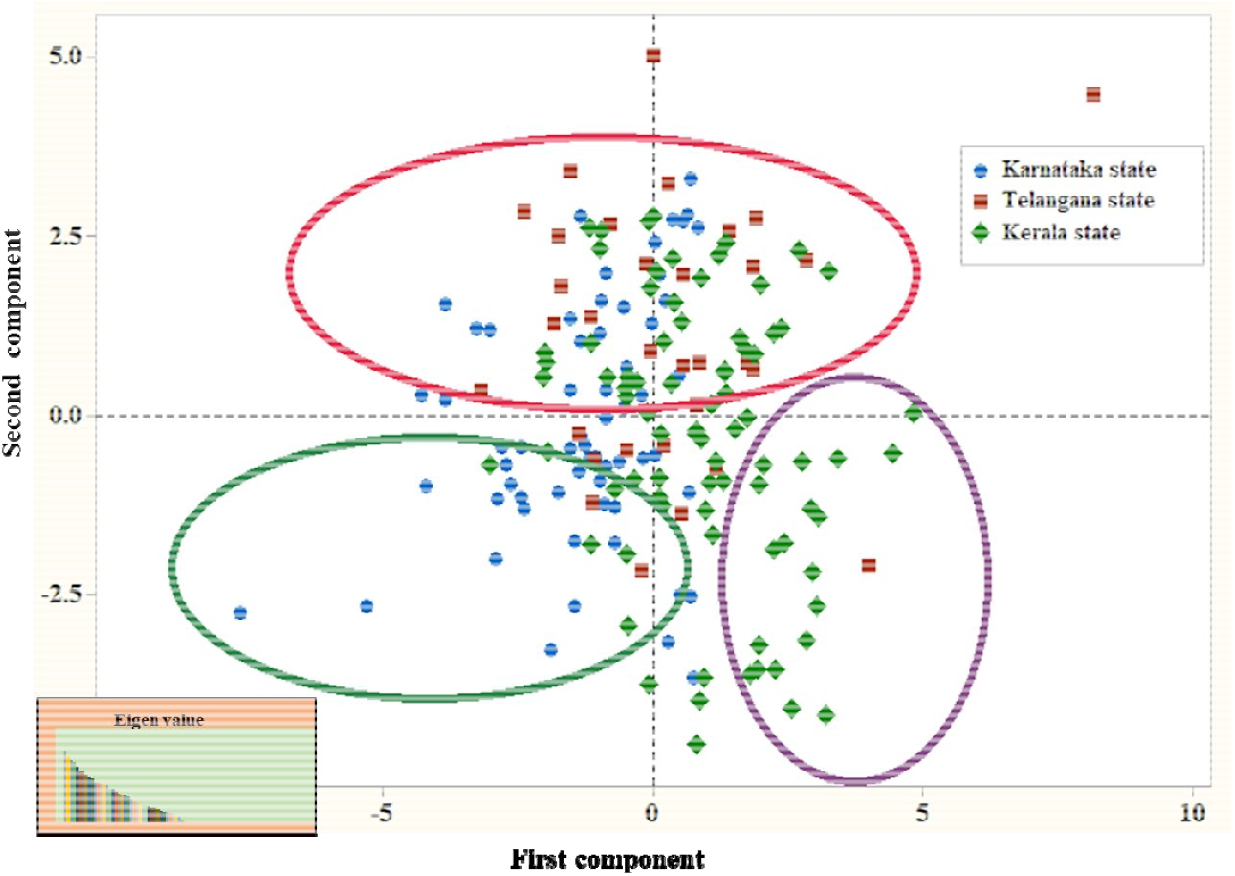
Principal Component Analysis of 177 accessions from 14 populations of *S. album* based on 39 microsatellite markers. The blue represents Karnataka genotypes, the green filled dots represent Kerala and the red filled dots represent Telangana state individuals. The first two Principle components explain percentage of total variation for PC1 6.0% and PC2 5.5% respectively

**Table 6.** Principal Component Analysis (PCA) of selected *S. album* accessions by using 39 genic SSR and genomic SSR markers

| **PC** | **Eigen value** | **Variation (%)** | **Cumulative (%)** |
| --- | --- | --- | --- |
| PC1 | 3.81 | 6.02 | 6.03 |
| PC2 | 3.52 | 5.53 | 11.67 |
| PC3 | 3.36 | 5.36 | 16.83 |
| PC4 | 3.23 | 5.18 | 22.07 |
| PC5 | 2.94 | 4.63 | 26.62 |
| PC6 | 2.73 | 4.41 | 30.99 |
| PC7 | 2.75 | 4.38 | 35.22 |
| PC8 | 2.56 | 4.02 | 39.29 |
| PC9 | 2.43 | 3.88 | 43.11 |
| PC10 | 2.31 | 3.71 | 46.97 |
| PC11 | 2.12 | 3.45 | 50.23 |
| PC12 | 2.17 | 3.37 | 53.57 |
| PC13 | 2.05 | 3.22 | 56.83 |
| PC14 | 2.01 | 3.25 | 59.98 |
| PC15 | 1.95 | 3.01 | 63.03 |
| PC16 | 1.79 | 2.77 | 65.79 |
| PC17 | 1.69 | 2.64 | 68.32 |
| PC18 | 1.63 | 2.59 | 70.98 |
| PC19 | 1.53 | 2.41 | 73.32 |
| PC20 | 1.45 | 2.35 | 75.68 |
| PC21 | 1.41 | 2.27 | 77.82 |
| PC22 | 1.34 | 2.13 | 79.96 |
| PC23 | 1.27 | 2.02 | 81.93 |
| PC24 | 1.15 | 1.81 | 83.75 |
| PC25 | 1.08 | 1.74 | 85.48 |
| PC26 | 0.93 | 1.55 | 86.86 |
| PC27 | 0.91 | 1.47 | 88.31 |
| PC28 | 0.88 | 1.48 | 89.62 |
| PC29 | 0.83 | 1.33 | 90.93 |
| PC30 | 0.77 | 1.21 | 92.24 |
| PC31 | 0.76 | 1.24 | 93.37 |
| PC32 | 0.70 | 1.15 | 94.49 |
| PC33 | 0.62 | 1.07 | 95.53 |
| PC34 | 0.57 | 0.95 | 96.32 |
| PC35 | 0.57 | 0.93 | 97.23 |
| PC36 | 0.50 | 0.87 | 98.06 |
| PC37 | 0.45 | 0.72 | 98.71 |
| PC38 | 0.34 | 0.57 | 99.35 |
| PC39 | 0.28 | 0.48 | 99.73 |

| PC | Eigen value | Variation (%) | Cumulative (%) |
| --- | --- | --- | --- |
| PC1 | 3.81 | 6.02 | 6.03 |
| PC2 | 3.52 | 5.53 | 11.67 |
| PC3 | 3.36 | 5.36 | 16.83 |
| PC4 | 3.23 | 5.18 | 22.07 |
| PC5 | 2.94 | 4.63 | 26.62 |
| PC6 | 2.73 | 4.41 | 30.99 |
| PC7 | 2.75 | 4.38 | 35.22 |
| PC8 | 2.56 | 4.02 | 39.29 |
| PC9 | 2.43 | 3.88 | 43.11 |
| PC10 | 2.31 | 3.71 | 46.97 |
| PC11 | 2.12 | 3.45 | 50.23 |
| PC12 | 2.17 | 3.37 | 53.57 |
| PC13 | 2.05 | 3.22 | 56.83 |
| PC14 | 2.01 | 3.25 | 59.98 |
| PC15 | 1.95 | 3.01 | 63.03 |
| PC16 | 1.79 | 2.77 | 65.79 |
| PC17 | 1.69 | 2.64 | 68.32 |
| PC18 | 1.63 | 2.59 | 70.98 |
| PC19 | 1.53 | 2.41 | 73.32 |
| PC20 | 1.45 | 2.35 | 75.68 |
| PC21 | 1.41 | 2.27 | 77.82 |
| PC22 | 1.34 | 2.13 | 79.96 |
| PC23 | 1.27 | 2.02 | 81.93 |
| PC24 | 1.15 | 1.81 | 83.75 |
| PC25 | 1.08 | 1.74 | 85.48 |
| PC26 | 0.93 | 1.55 | 86.86 |
| PC27 | 0.91 | 1.47 | 88.31 |
| PC28 | 0.88 | 1.48 | 89.62 |
| PC29 | 0.83 | 1.33 | 90.93 |
| PC30 | 0.77 | 1.21 | 92.24 |
| PC31 | 0.76 | 1.24 | 93.37 |
| PC32 | 0.70 | 1.15 | 94.49 |
| PC33 | 0.62 | 1.07 | 95.53 |
| PC34 | 0.57 | 0.95 | 96.32 |
| PC35 | 0.57 | 0.93 | 97.23 |
| PC36 | 0.50 | 0.87 | 98.06 |
| PC37 | 0.45 | 0.72 | 98.71 |
| PC38 | 0.34 | 0.57 | 99.35 |
| PC39 | 0.28 | 0.48 | 99.73 |

## 4. Discussion

A number of causes have been endorsed to the depletion of *S. album* populations specifically due to overharvesting and illegal poaching in natural habitats (Azeez et al., 2009; Naseer et al., 2012). Therefore conservation and systematic cultivation in terms of identifying the elite tree/populations of *S. album*, genetic diversity study is essential as it provides the important factors that direct the variability (adaptation, migration), inbreeding and gene flow (Naseer et al., 2012; Patel et al., 2016).

### 4.1. Genetic Diversity and variation

In the current study we estimated the genetic diversity parameters of 177 *S. alum* accessions at 39 loci. Overall genetic diversity among the populations was high (Na=9.39 and He 0.85) (Table 2). Similar results were obtained in previous studies of *Santalum* and other tree species like *O. lanceolata* (Otieno et al., 2016), *S. spicatum* (Millar et al., 2012), *S. lanceolatum* (Jones et al., 2010), *S. leptocladum* (Jones et al., 2010), *S. album* (Fatima et al., 2019b) and *Phyllostachys violascens*, (Cai et al., 2019). In contrast to our study, low Na was observed in *S. album* by using genic SSR markers (Fatima et al., 2019a) and, *Ambrosia artemisiifolia*, (Meyer et al., 2017). The PIC value of 14 populations ranged from 0.83-0.96 (Table 2) in which 10 populations were highly polymorphic (PIC>0.8). The average PIC (0.85) was higher than that of previous study on *S. album, Jatropha curcas*, (Naseer et al., 2012; Wen et al., 2014). The PIC value is changed by number of genetic markers, types of SSR markers, the SSR motif repeats and the analysis method (Liu et al., 2017; Chen et al., 2020). As in another study high level of polymorphism was observed in *S. album, Populus tomentosa, Quercus variabilis* and *Eucalyptus tereticornis* respectively (Patel et al., 2016; Du et al., 2012; Shi et al., 2017; Zhijiao et al., 2016).

### 4.2. Population structure

Investigation of population structure using several approaches supported the evolutionary divergence of *S. album* populations in selected states. In this study we analyzed the population structure of *S. album* at different levels. At the individual level, cluster, structure and PCA analysis were performed on 177 accessions which divided them into groups. The Bayesian structure analysis and PCA of *S. album* defined four groups and provided scattered genetic divergence among the populations from the selected three states *viz*, Karnataka, Kerala and Telangana (Fig. 2 and Fig. 6). The AMOVA revealed these four groups were not differentiated among the populations (3% P<0.001) (Table 3). The structure result also confirmed AMOVA test and showed presence of genetic diversity within each presumed population groups. Similar results were reported in *S. album* and *Corylus mandshurica* (Fatima et al., 2019a; Fatima et al., 2019b; Zong et al., 2015;). The obtained results indicated the mix lineage of the nuclear genome and distributed randomly among the populations. Within *S. album* populations distributed in southern India, no clear geographical genetic differentiation was found by the NJ tree, Bayesian and the PCA analysis (Fig. 1, Fig. 2 and Fig. 6). In contrast to present study genotypes of similar populations/geographical region were clustered together as per their geographical locations *viz. Tectona grandis, Cabralea, Canjerana and Swietenia macrophylla* (Alcantara et al., 2013; Sreekanth et al., 2012; Fofana, et al., 2009; Melo et al., 2014; Alcala et al., 2014; Lemes et al., 2002). UPGMA cluster analysis showed IWC L1 KA and Nel L2 KA populations were clustered together in cluster II and confirmed the seedling source of *S. album*. Similar result was observed in another study of *S. album* by oil specific genic markers (Fatima et al., 2019a). On the other hand a previous study using random genomic markers indicated mixed clustering of *S. album* populations (Fatima et al., 2019b). This might revealed the result based on the genic markers transferability within the species, high reproducibility and their conserved nature (Powell et al., 2007; Muriira et al., 2018).

## 5. Conclusion

In the present study, the genetic diversity and population structure of *S. album* genotypes were analyzed. The result revealed the high genetic variability and genetic differentiation in fourteen populations of *S. album* of three selected states of India. The observed variation in selected populations of *S. album* will aid in the selection of elite genotypes for genetic improvement, conservation management and tree breeding program. Additionally, these markers would be a useful tool for investigating genetic diversity, genetic structure of other states natural populations and plantations of *S. album*. The information obtained by both genomic and genic SSR markers offer precise mark on the distribution of the genetic diversity among the selected populations.

## Acknowledgement

Authors are thankful to the Director, Group Co-ordinator Research, Head-Genetics and Tree Improvement Division, Dr. D. Annapurna GTI Division (PDF), Institute of Wood Science and Technology for encouragement to carry out the present study.

## Author contributions

First author design the experiment, completed laboratory work and written manuscript. All other authors reviewed the manuscript and helped in formatting.

## Availability of data and materials

We declare that all data generated or analyzed during this study are included in this manuscript.

## Highlights

- *Santalum album* L. is important aromatic tropical tree species.
- In the present study, 39 genic and genomic SSR markers were used to analyze genetic diversity and population structure of 177 *S. album* from 14 populations of three states.
- The result revealed the high genetic variability and genetic differentiation in fourteen populations of *S. album* from the selected states.
- The genetic diversity information of *S. album* populations could be used for selection of superior genotypes to promote the genetic tree improvement of *S. album*.

---

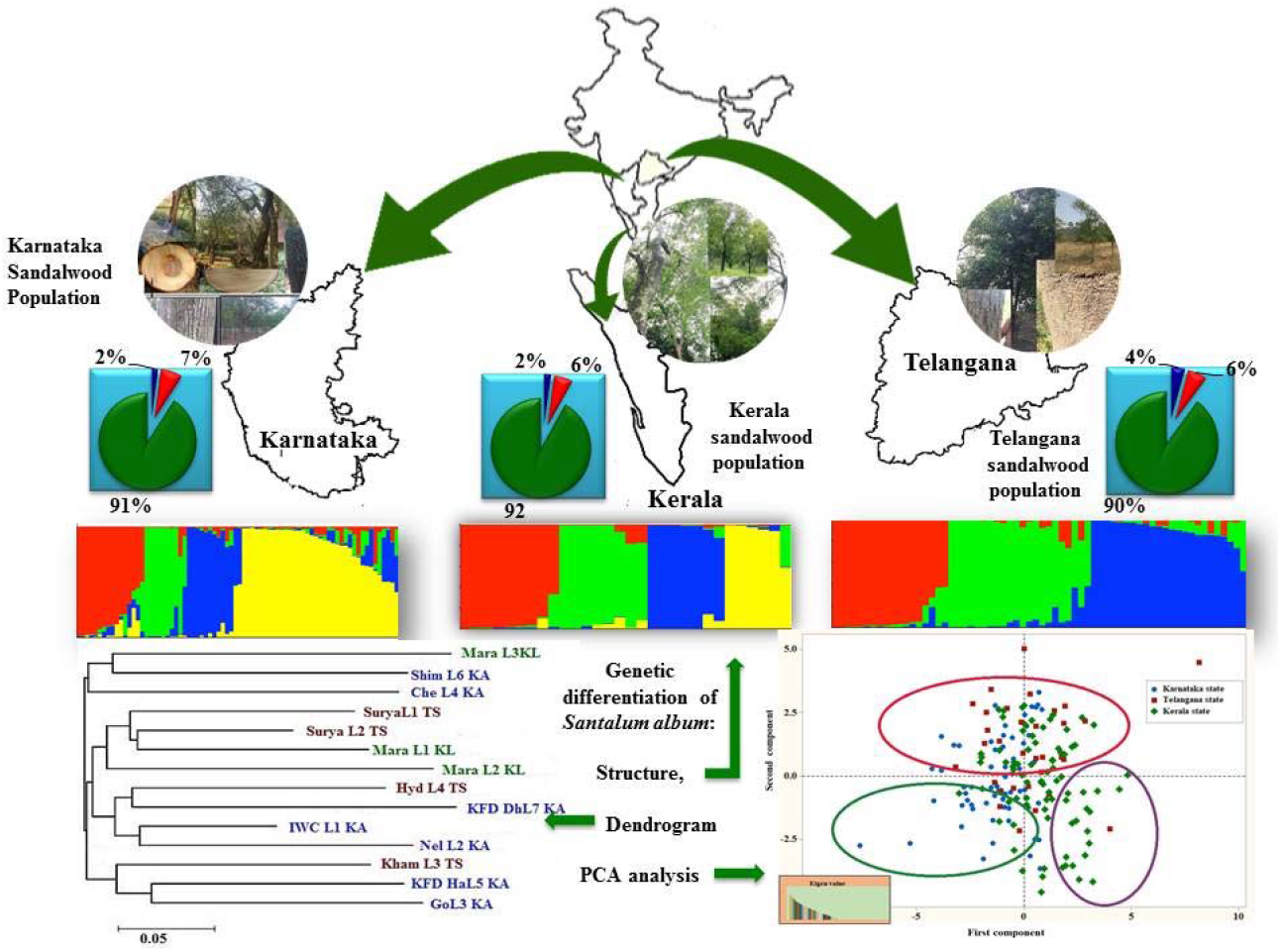

